# Clinically Important sex differences in GBM biology revealed by analysis of male and female imaging, transcriptome and survival data

**DOI:** 10.1101/232744

**Authors:** Wei Yang, Nicole M. Warrington, Sara J. Taylor, Eduardo Carrasco, Kyle W. Singleton, Ningying Wu, Justin D. Lathia, Michael E. Berens, Albert H. Kim, Jill S. Barnholtz-Sloan, Kristin R. Swanson, Jingqin Luo, Joshua B Rubin

## Abstract

Sex differences in the incidence and outcome of human disease are broadly recognized but in most cases not adequately understood to enable sex-specific approaches to treatment. Glioblastoma (GBM), the most common malignant brain tumor, provides a case in point. Despite well-established differences in incidence, and emerging indications of differences in outcome, there are few insights that distinguish male and female GBM at the molecular level, or allow specific targeting of these biological differences. Here, using a quantitative imaging-based measure of response, we found that temozolomide chemotherapy is more effective in female compared to male GBM patients. We then applied a novel computational algorithm to linked GBM transcriptome and outcome data, and identified novel sex-specific molecular subtypes of GBM in which cell cycle and integrin signaling were identified as the critical determinants of survival for male and female patients, respectively. The clinical utility of cell cycle and integrin signaling pathway signatures was further established through correlations between gene expression and *in vitro* chemotherapy sensitivity in a panel of male and female patient-derived GBM cell lines. Together these results suggest that greater precision in GBM molecular subtyping can be achieved through sex-specific analyses, and that improved outcome for all patients might be accomplished via tailoring treatment to sex differences in molecular mechanisms.

**One Sentence Summary:** Male and female glioblastoma are biologically distinct and maximal chances for cure may require sex-specific approaches to treatment.

## Introduction

Current epidemiological data indicate that significant sex differences exist in the incidence of cardiovascular disease, disorders of the immune system, depression, addiction, asthma and cancers (*1*-*4*), including glioblastoma (GBM) (*5*). While sex differences in disease incidence and severity may parallel variation in circulating sex hormone levels, in many cases, sex differences exist across all stages of life, indicating independence from acute hormone action (*3*, *6*). Sex differences in GBM are evident in all age groups, and therefore cannot be solely the consequence of activational effects of sex hormones (*5*, *7*-*11*). Enumerating the molecular bases for sex differences in GBM is likely to reveal fundamental modulators of cancer risk and outcome, as well as guide sex-specific components of precision medicine approaches to cancer treatment.

Identifying the basis for sex differences in cancer biology cannot be accomplished by analysis of merged male and female datasets. Instead, it requires comparison of results from parallel analyses of male and female data. The importance of this was recently highlighted in a study of asthma, a disease driven by both genetic and environmental factors, which occurs in twice as many boys as girls. Mersha *et al*. examined the influence of genetic variants on asthma, including an analysis of shared and sex-specific variant effects (*2*). Of 47 variants that correlated with asthma risk in the sex-specific analyses, only 21 were detected in the combined analysis, suggesting that biologically important mechanisms of disease were obscured by a “net cancelling effect” that arose from opposing effects of the interactions between sex and genetic variation. Moreover, even when males and females exhibit similar characteristics of a disease, the mechanisms driving the disease state may differ (*2*, *12*, *13*). For example, despite equal tumor incidence in males and females, polymorphisms in *AC8* in patients with NF1 elevate risk of low-grade glioma in female patients while reducing the risk in male patients (*14*).

While low-grade glioma incidence is nearly identical in males and females, malignant brain tumors in general occur more commonly in males, regardless of patient age or geographical location (*6*, *15*) (*5*, *11*). From multiple recent reports, GBM occurs with a male to female ratio of 1.6:1 (*5*, *8*-*10*). More specifically, while concepts of molecular subtypes of GBM are still evolving (*16*), of the four originally described transcriptional subtypes of GBM, three – Mesenchymal, Proneural and Neural GBM – exhibit a 2:1 male to female incidence ratio, while Classical GBM occurs with equal incidence (*17*, *18*). To date, analyses of the transcriptome data from which these molecular subtypes were derived have been performed with merged male-female data and have not yielded new insights into the molecular basis for sex differences in GBM incidence.

In addition to sex differences in incidence, there are emerging analyses that suggest outcome from GBM may also differ between males and females. In a study analyzing more than 27,000 patients, Trifiletti *et al*. found that female sex was associated with longer survival (*10*). Similarly, female patients exhibited longer survival from gliosarcoma (*8*), and being female was associated with better outcome in a newly developed nomogram for predicting GBM patient survival (*9*). Thus, the elucidation of mechanisms for sex differences in the development and treatment of GBM has a substantial potential to improve outcome for all patients by refining our understanding of disease causation and outcome.

In the current study, we performed quantitative analyses of temozolomide response in male and female GBM patients using a validated magnetic resonance imaging-based algorithm for calculating tumor growth velocities. We also applied a novel computational algorithm to male and female GBM transcriptome data in order to gain new insights into the significance and biological basis of sex differences in GBM. Our studies indicate that temozolomide is more effective for females than for males with GBM and that, for current standard of care treatment with surgery, radiation and temozolomide, survival in males is determined by expression of cell cycle regulators, while in females it is determined by expression of integrin signaling pathway components. These studies provide a novel and coherent view of sex differences in GBM biology and their clinical ramifications. They strongly endorse the development of diagnostics and treatments that incorporate sex differences in GBM biology.

## Results

### Temozolomide treatment is more effective in female compared to male GBM patients

Sex differences in GBM incidence have been repeatedly reported (*5*, *7*-*11*). Moreover, several recent studies have suggested that being female is associated with better outcome from GBM in both adults and children (*8*-*10*, *19*). The introduction of temozolomide as a component of tri-modal care for adults with GBM has improved outcomes somewhat and highlighted factors, like MGMT promoter methylation, that significantly impact on response and survival (*20*, *21*). Thus, we wondered whether sex differences in GBM survival are a consequence of differential temozolomide effects on males versus female patients. To answer this question we utilized a magnetic resonance (MR) imaging data-based analysis, with which the velocity of radial tumor expansion can be derived (*22*-*25*). This parameter, which has been prospectively validated to correlate with outcome (*26*, *27*), was measured approximately every two months in a cohort of 111 GBM patients treated with standard-of-care surgery, focal irradiation (RT) and systemic temozolomide (TMZ) chemotherapy (*20*, *21*). Analysis of these serial MR images indicated that female patients exhibited a greater response to TMZ than male patients. This was evident as a steady decline in growth velocity during TMZ treatment for female patients; a change that was not detected in male patients (**Figure 1A**). The velocity of radial expansion was significantly different between males and females by the 3^rd^ interval scan, which typically bridged the 5^th^ and 6^th^ adjuvant cycles of TMZ (**Figure 1B**, p=0.01951). These data suggest that females with GBM may benefit more, albeit transiently, from TMZ, than males with GBM.

**Figure 1:**
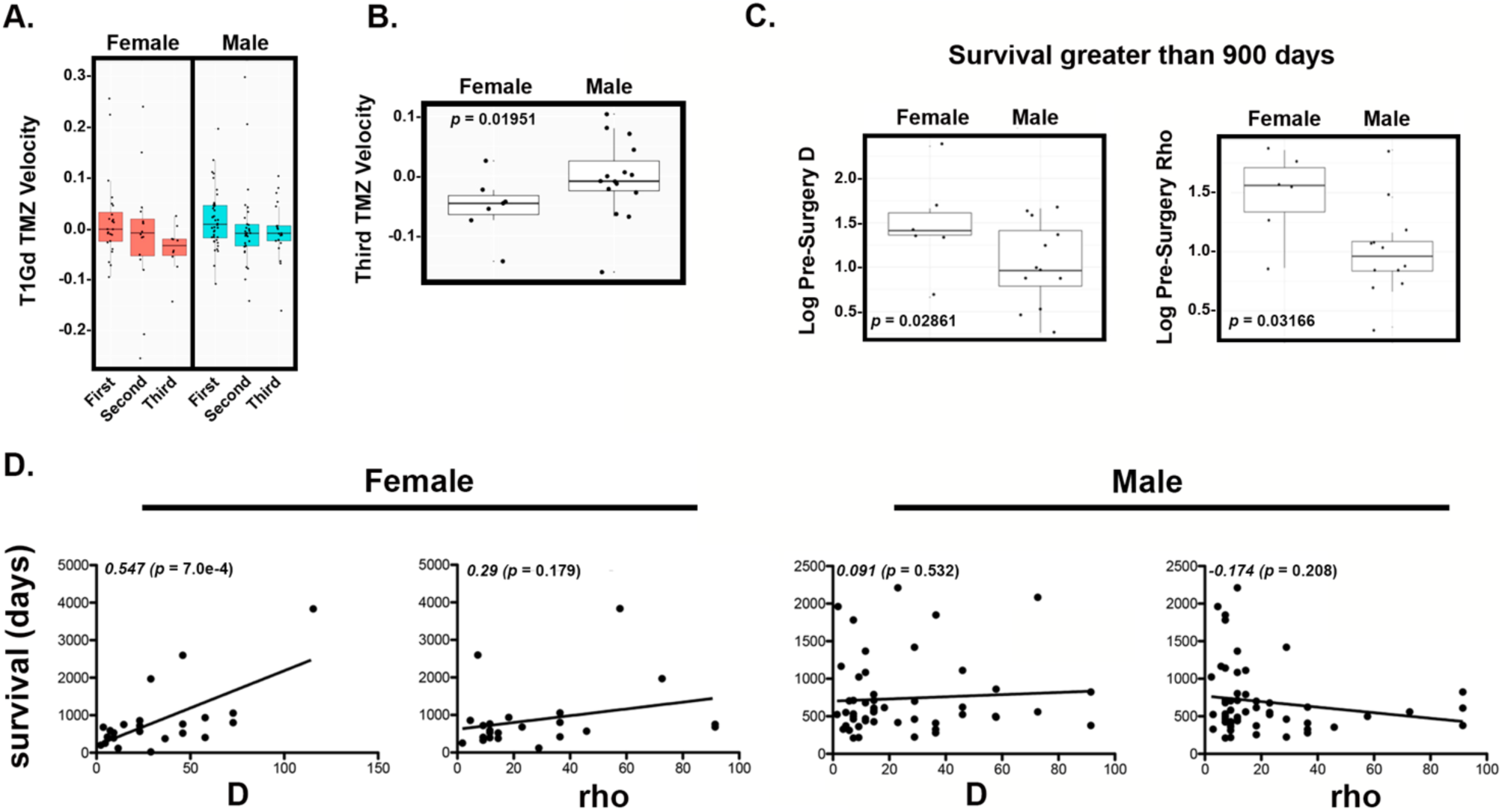
Sex differences in MRI-based metrics of therapeutic responses and their correlation with survival. **(A)** Tumor growth velocities calculated from serial MRI images exhibit progressive decline for female but not male patients treated with temozolomide (TMZ). **(B)** After six cycles of TMZ (third velocity) females exhibit statistically significant differences in tumor growth velocity (p = 0.01951) as determined by Chi-squared analysis. **(C)** Pretreatment D (p = 0.02861) and ρ (p = 0.03166) were significantly greater in female GBM patients who survived greater than 900 days as compared to males with similar survival.ρ value was calculated using a Welch Two Sample *t*-test. (D) Sex-specific correlations between D, ρ and survival were determined. Best-fit lines are presented for each dataset along with Spearman correlation coefficients and the associated p-values.

To further investigate the basis for this difference in response, we applied an established mathematical model of glioma proliferation and invasion (*24*, *25*) to each patient’s pre-surgical MRIs (T1-gadolinium and T2 sequences), in order to estimate patient-specific tumor net proliferation (rho (ρ), 1/year) and net infiltration rates (D, mm^2^/yr) (*23*, *25*, *28*). Overall, those females (6, 16%) with the longest survival (> 900 days) exhibited significantly greater values for D (chi-square, p=0.02861) and ρ (chi-square, p=0.03166) compared to long-lived male patients (12, 16%) (**Figure 1C**). We next asked whether D and ρ would be predictive of outcome within the male and female patient populations and distinguish longer-lived from shorter-lived males or females. We found a significant positive correlation between D and survival in female (p=7.0e-4) but not male (p=0.532) patients (**Figure 1D**). While neither male nor female survival was significantly correlated with values for ρ, long-lived males had significantly lower values for rho (p=0.03166 compared to females **Figure 1C**), and there was a trend towards a negative correlation between ρ and survival for males (p=0.208, **Figure 1D**). This was in contrast to the trend towards positive correlation between ρ and survival in females (p=0.179, **Figure 1D**). These data suggest that males and females with GBM exhibit different acute responses to temozolomide and that this may contribute to the differences in their outcome. Together with the established sex differences in incidence, these data suggest that the biology of male and female GBM may be distinct and that outcomes for all patients might be improved if therapy were better tailored to patient sex.

### Sex differences in GBM biology are revealed by JIVE decomposition

Using JIVE to integratively decompose the male and the female transcriptome data into three orthogonal components, we identified the joint structure that was common to both sexes, the individual structure that was specific to each sex, and additionally, the residuals (**Supplemental Figure 1**). We focused on the male/female-specific components to examine subtypes within each sex separately. The sex specific expression components underwent independent hierarchical clustering to identify patient subgroups and five male (mc1-5) and five female (fc1-5) clusters were identified (**Supplemental Figure 2, Supplemental Table 1**). The heat maps of the male joint structure across the male GBM patients and the female joint structure across the female GBM patients indicated that the joint structures extracted by JIVE closely captured the dominant molecular signatures defining the TCGA GBM subtypes (**Supplemental Figure 3**). However, the joint component only explained ~45% of the total variance in the transcriptomes for each sex, while the sex-specific components, independent of the joint components, explained a great proportion of the remaining variability. Specifically, the male-specific component accounted for 38.5% of the total variability in the male transcriptome, and the female-specific component explained 33.6% of the total variability in the female transcriptome (**Supplemental Figure 4**).

Cases from multiple TCGA molecular subtypes (*18*) were distributed to each of the male/female clusters, indicating successful separation of the individual components from the joint structure components and increasing the likelihood that this approach could reveal sex effects on gliomagenic mechanisms. The one exception was fc3, 70% of which were Proneural subtype tumors with IDH1 mutation (7 IDH1 mutants, 3 WT, **Supplemental Table 2**). In contrast, male Proneural subtype tumors with IDH1 mutation were distributed across three of the new male clusters.

The extracted male and female individual components exhibited distinct patterns compared to their counterpart joint structure, and more importantly, the male-specific component showed distinct patterns compared to the female-specific component (Figure 2). We hypothesized that focused analyses on the extracted sex-specific components would reveal which gliomagenic mechanisms are most characteristic of male versus female cases.

**Figure 2:**
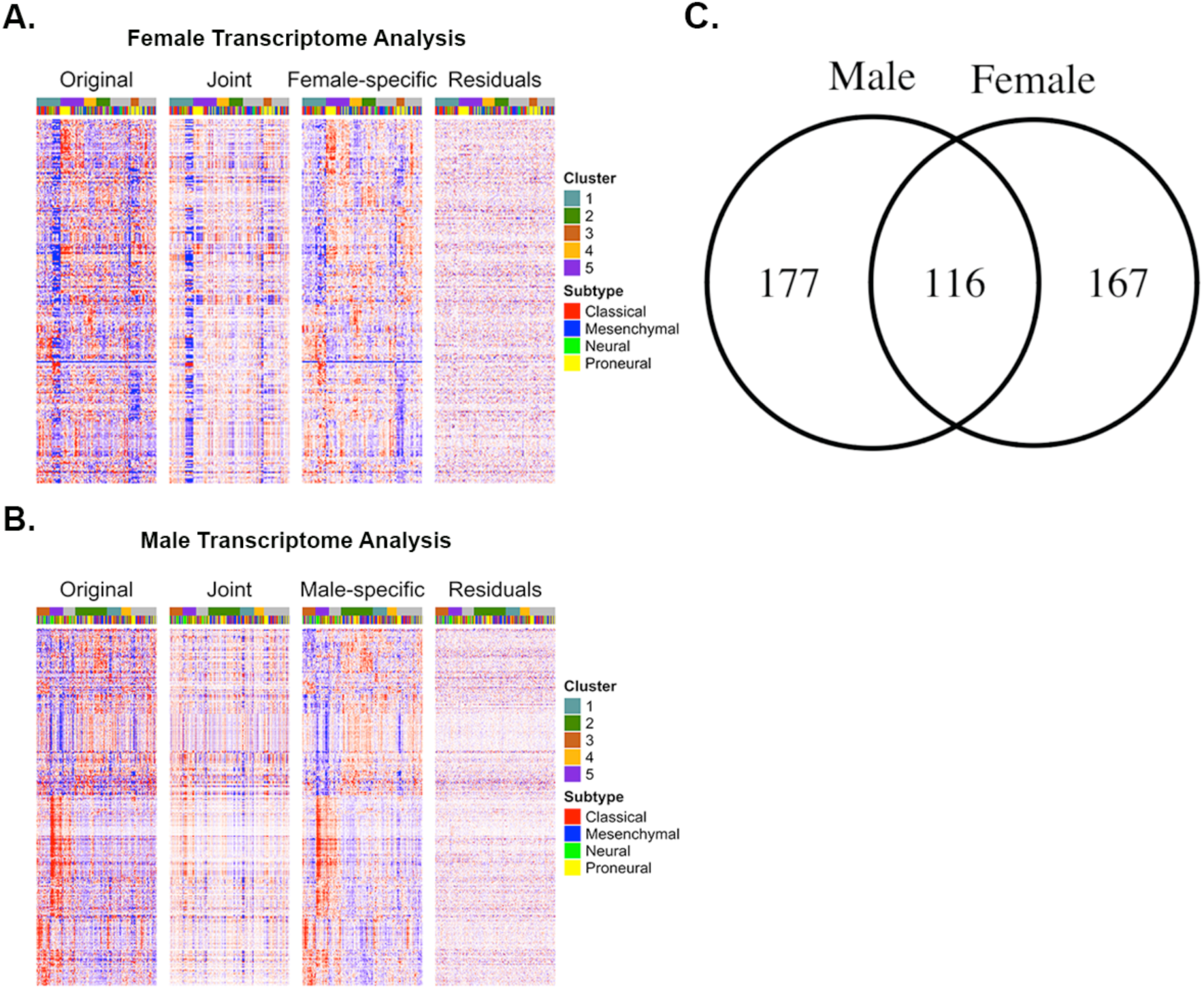
Heatmaps of Joint and Sex Specific Expression Components of TCGA GBM transcriptome data revealed by JIVE. The heatmaps visualize each expression component. Each row represents a gene and each column a patient sample. For each patient, there are two color codes presented above the heatmap. These identify their assignment to sex-specific clusters and to TCGA molecular subtypes. Samples were ordered by sex-specific clusters. The original female **(A)** and male **(B)** expression data were decomposed into the shared expression component common to both sexes (“Joint”) and the expression component individual to each sex (“Female-specific” and “Male-specific”) and residuals as indicated. The female-relevant heatmaps **(A)** show 283 signature genes that define the five female-specific clusters and the male-relevant heatmaps **(B)** show 293 signature genes that define the five male-specific clusters. **(C)** Venn diagram of male and female signature genes indicating 116 genes are in common.

### Survival differences exist among sex-specific clusters

To establish the importance of the sex-specific clusters, we next determined whether the sex-specific clusters in the TCGA data were associated with differences in survival outcomes. Kaplan-Meier (KM) analyses of male/female clusters confirmed that survival differences exist among both male and female clusters. Not surprisingly, fc3, in which 70% of the cases are IDH1 mutant, exhibited significantly better disease free survival (DFS) with a median time to progression (TTP) of 1758 days compared to each of the other four female clusters (fc1 259 days, *p*=3.3e-5; fc2 289 days, *p*=5e-4; fc4 182 days, *p*=1.64e-4; fc5 350 days, *p*=9.6e-5, **Figure 3A,** **Supplemental Table 3**). In contrast, while IDH1 mutant cases segregated nearly equally to mc2, 3 and 5, only mc3 (median TTP 408 days) and mc5 (median TTP 262 days) were associated with prolonged DFS compared to other male clusters (mc1 240 days (*p*=1.2e-2 (mc3)), mc2 186 days (*p*=7.1e-3 (mc3), *p*=2.8e-2 (mc5)); mc4 158 days (*p*=7.3e-3 (mc3), *p*=1.6e-2 (mc5)), **Figure 3B,** **Supplemental Table 3**), suggesting greater phenotypic divergence among male IDH1 mutant GBMs. Similar results were observed in female but not male OS in the TCGA dataset (**Supplemental Table 3, Supplemental Figure 5**). Sensitivity analyses were conducted on survival results after removing IDH1 mutant samples in the female and male clusters. Only three cases in fc3 were IDH wildtype, but remarkably, all of them were alive at 5 years (median survival for fc3 was not calculable; fc1 259 days (*p*=1.6e-2); fc2 322 days (*p*=4.3e-2); fc4 274 days (*p*=4.9e-2); fc5 350 days (*p*=1.2e-2), **Figure 3C,** **Supplemental Table 3**). Similarly, the survival benefit of mc3 and mc5 remained intact after removal of the IDH1 mutant cases (mc3 426 days; mc5 350 days; mc1 204 days ((*p*=2.24e-4 vs. mc3), (*p*=2.6e-2 vs. mc5)); mc2 176 days ((*p*=8e-4 vs. mc3), (*p*=1.4e-2 vs. mc5)); mc4 131 days ((*p*=1.3e-2 vs. mc3), (*p*=4.3e-2 vs. mc5)), **Figure 3D,** **Supplemental Table 3**). These results suggest that the survival effects of fc3, mc3, and mc5 are independent of IDH1 mutational status.

**Figure 3:**
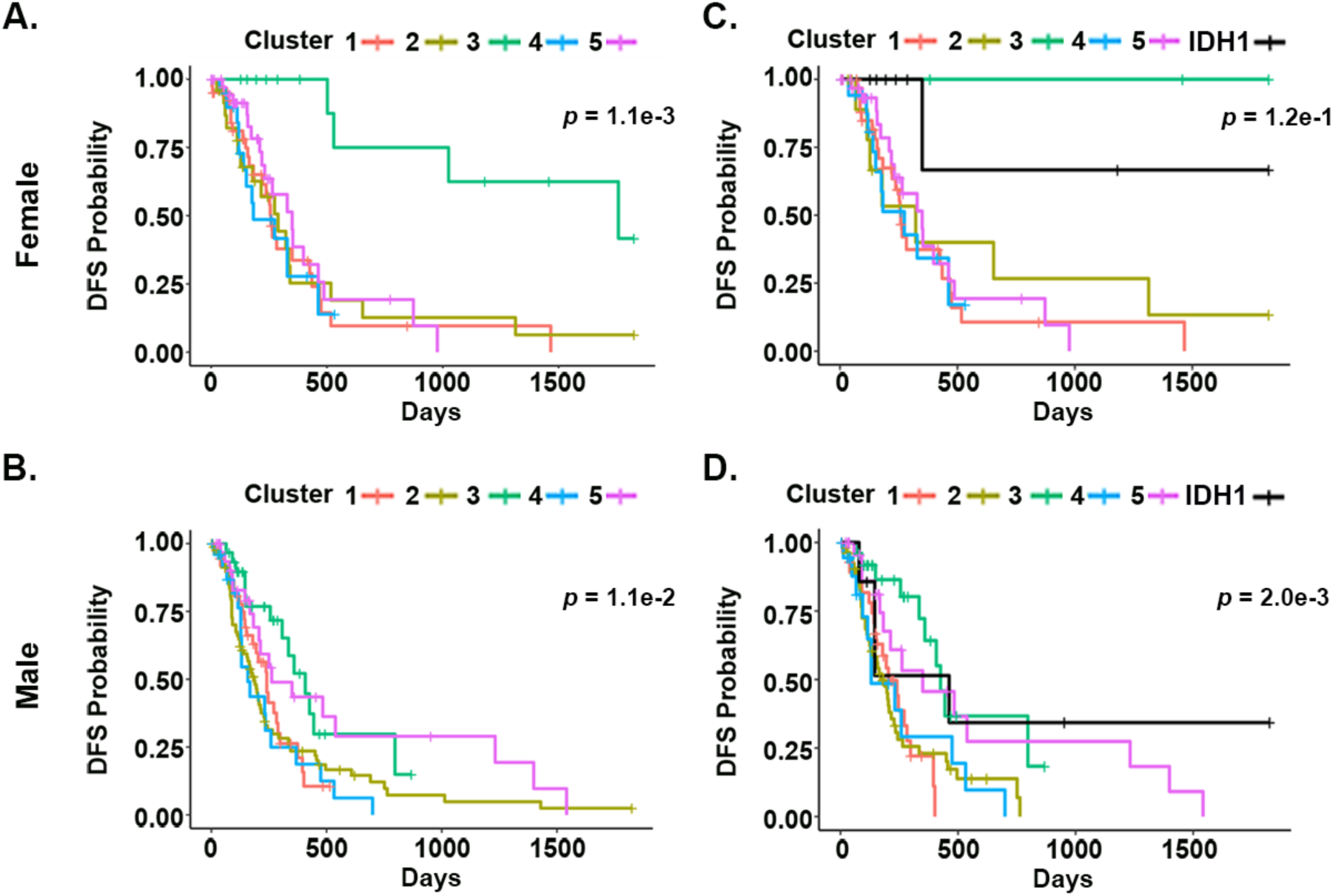
Disease Free Survival of Sex-Specific Clusters in TCGA GBM dataset. **(A)** Disease free survival (DFS) in TCGA-derived female clusters (1-5). Survival in cluster 3 is significantly greater than each of the other clusters. **(B)** Disease free survival (DFS) in TCGA-derived male clusters (1-5). Survival in clusters 3 and 5 are significantly greater than each of the other clusters. **(C)** Disease free survival (DFS) in TCGA-derived female clusters (1-5) after removal of IDH1 mutant cases. IDH1 mutant cases are plotted as an independent cluster. A survival benefit of cluster 3 remains after the removal of IDH1 mutant cases. **(D)** Disease free survival (DFS) in TCGA-derived male clusters (1-5) after removal of IDH1 mutant cases. IDH1 mutant cases are plotted as an independent cluster. A survival benefit of clusters 3 and 5 remains after the removal of IDH1 mutant cases. Overall log rank test*p* value is shown comparing across all the groups presented in each panel (see **text** and **Supplemental Table 3** for the *p*-values and hazard ratios for all pairwise comparisons).

### Validation analyses

In order to validate the male and female cluster-specific survival profiles, transcriptome data of GSE13041 and GSE16011 were decomposed with the JIVE principal components (PCs) from the TCGA data analysis (**Supplemental Figure 1**). Only overall survival was evaluable in these datasets, and therefore we were limited to an analysis of this survival endpoint. An overall survival benefit of fc3 and mc5 was validated in these datasets (**Supplemental Table 3**). In an integrated analysis of all the three datasets, median survival for fc3 was 1179 days and compared to 396 days for fc1 (*p*=1.5e-4), 486 days for fc2 (*p*=2.6e-6), 423 days for fc4 (*p*=3.4e-8) and 310 days for fc5 (*p*=5.0e-7) (**Figure 4A,** **Supplemental Table 3**). Median survival for mc5 was 675 days and compared to 413 days for mc1 (*p*=6.9e-6), 360 days for mc2 (*p*=6.6e-8), 394 days for mc3 (*p*=8.7e-4), 323 days for mc4 (*p*=2.5e-5). Of the two validation datasets, only GSE16011 specified IDH mutational status. In this dataset, IDH1 mutant tumors were disproportionately distributed to fc3 and more broadly to multiple male clusters (**Supplemental Table 2**).

**Figure 4:**
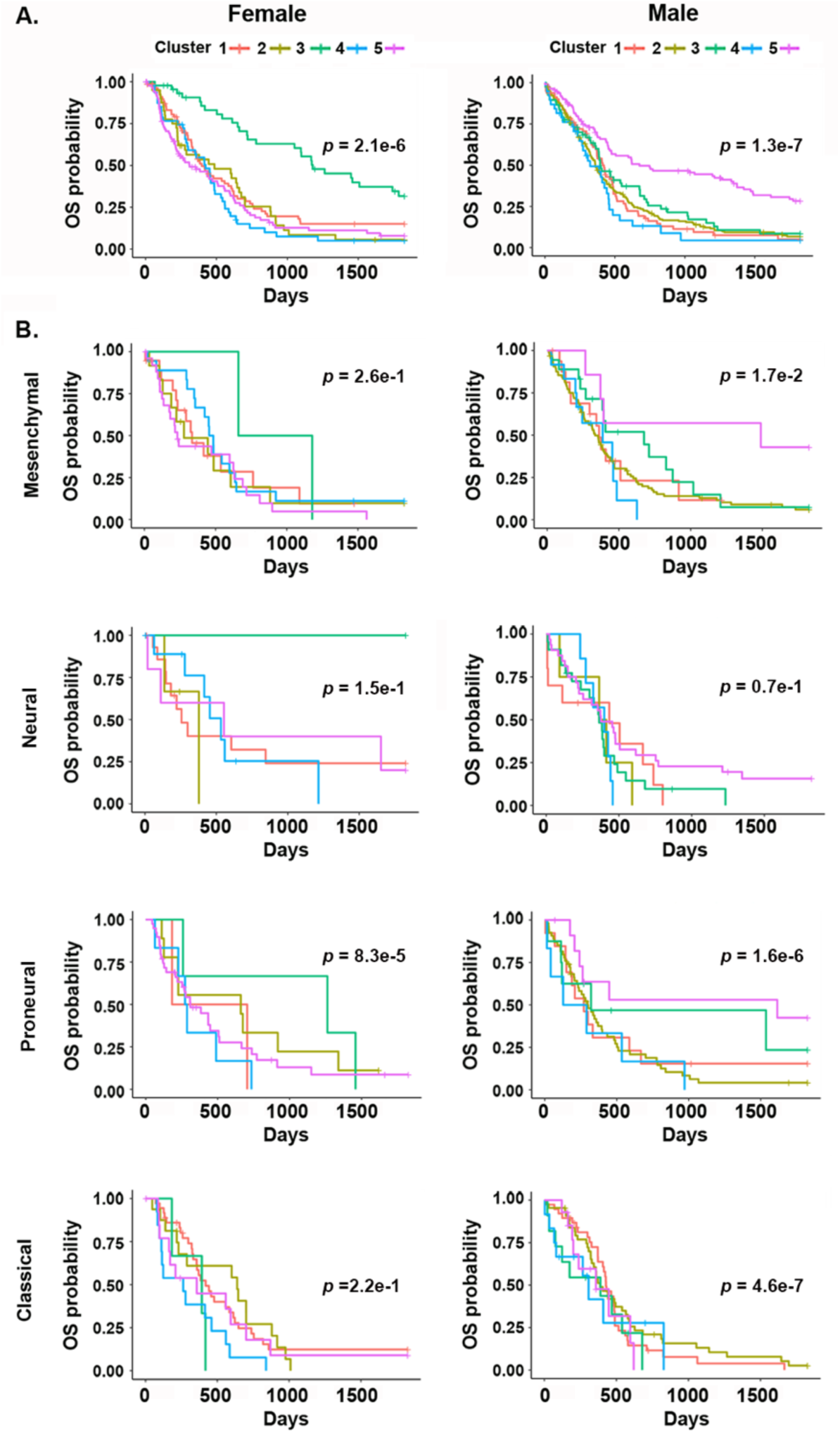
Sex-specific cluster survival benefits in merged TCGA, GSE16011 AND GSE13041 dataset. **(A)** Samples from GSE16011 and GSE13041 were assigned to male and female specific clusters and overall survival was plotted for each cluster in the combined TCGA, GSE16011 and GSE13041 dataset. Patients belonging to fc3 or mc5 exhibit significantly greater overall survival compared to all other clusters. Samples from all three datasets were assigned to TCGA subtypes and the sex-specific cluster effect on survival was evaluated. **(B)** Fc3 and mc5 exhibit a survival benefit for Mesenchymal, Neural and Proneural subtype tumors but not for Classical subtype tumors. Overall log rank test *p* values are shown comparing across all the groups presented in each panel (see **text** and **Supplemental Table 3** for the *p*-values and hazard ratios for all pairwise comparisons).

To gain further insights into cluster-specific effects on survival, we compared the survival differences of the male and female specific clusters within each Verhaak subtype. We found a consistent cluster effect in which Neural, Mesenchymal and Proneural specimens in mc5 and fc3 exhibited better survival than tumors of these same Verhaak subtypes that had clustered to mc1-4 or fc1,2, 4, or 5 (**Figure 4B**). Neither male nor female cluster effects were evident for the Classical subtype tumors, the only subtype for which there is no sex difference in incidence. These new data suggest that for those molecular subtypes of GBM in which sex impacts tumor incidence, sex also impacts patient survival. In addition, these findings indicate that sex can modulate the impact of specific gliomagenic mechanisms on survival, but that not all mechanisms, such as those underlying Classical subtype tumors, will be sensitive to the effects of sex. Together, these results suggest greater prognostic precision might be achieved through sex-specific molecular subtyping.

### Pathway Analysis

The survival advantage of fc3 and mc5 was evident regardless of whether the component tumors belonged to the Mesenchymal, Proneural, or Neural subtypes. This was in contrast to tumors belonging to the Classical subtype in which survival was unaffected by patient sex. The unequal effect of sex on survival for these different molecular subtype tumors suggests that the effects of sex are not mediated solely by factors like sex hormones, whose actions would be predicted to distribute equivalently across patients of a given sex regardless of their molecular subtype. Instead these findings indicate that either tumor cell intrinsic sex differences or an interaction between tumor intrinsic and microenvironmental sex differences determine responsiveness to treatment and patient survival. To gain insight into possible mechanisms underlying sex-specific survival benefits, we compared the survival and transcriptome expression of fc3 to mc5.

A substantial trend, though statistically not significant (*p*=0.213), towards longer survival was evident for fc3 (median survival 1179 days) compared to mc5 (median survival 675 days) based on KM analysis (**Figure 5A**). To test whether similar or distinct mechanisms accounted for these sex differences in outcome, we asked what distinguished fc3 and mc5 from the other female and male clusters, respectively. One hundred ninety-seven transcripts distinguished mc5 from the other male clusters, and 123 transcripts distinguished fc3 from the other female clusters (**Supplemental Tables 1 and 4**). Forty-three transcripts were shared between mc5 and fc3. Using the Genomatix Suite for pathway analysis, we found that the most significant pathways shared by the best surviving male and female tumors involved calcium/calmodulin signaling, including a large number of synaptic and other neuronal function genes (*SYN1*, *SYT1*, *SNAP25*, *HTR2A*, *CAMK2B*, *CAMK2G*, *NRGN*, *INA*, *NEFL*, *PCP4*) (**Figure 5B**). Examination of the female-specific transcripts revealed the Integrin signaling pathway as the most significant pathway that distinguished fc3 from other female clusters (**Figure 5C**). Six of the nine transcripts from this pathway (*PLAT*(*29*), *CHL1* (*30*, *31*), *FERMT1* (*32*), *PCDH8* (*33*), *IGFBP2* (*34*, *35*), *POSTN* (*36*)) have previously reported roles in glioma, and three (*PLAT*, I*GFBP2* and *POSTN*) have been reported to distinguish Proneural from Classical subtype GBM consistent with the high rate of IDH1 mutant tumors in this cluster. Six of the nine genes (*AK5*, *AMIGO2*, *PLAT*, *CHL1*, *PCDH8*, *IGFBP2*) were downregulated in fc3 compared to other female clusters suggesting that better survival in fc3 patients is favored by tumors with reduced integrin signaling (data not shown).

**Figure 5:**
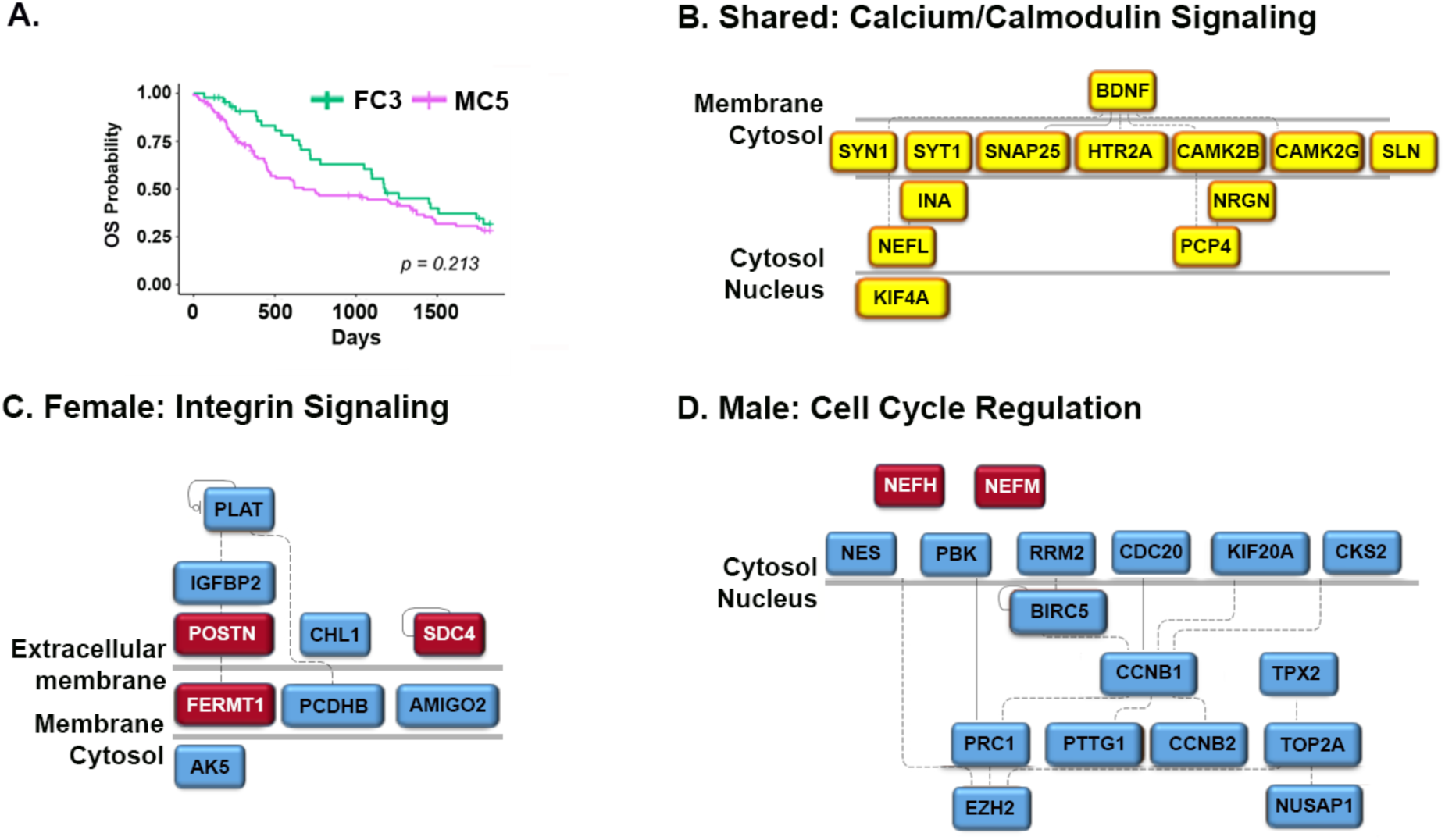
Analysis of genes and pathways that mediate better survival. **(A)** In the combined dataset, females assigned to female cluster 3 trend towards enhanced survival (median survival 1179 days) compared to males assigned to male cluster 5 (median survival 675 days). Genes that distinguished female cluster 3 and male cluster 5 from other female and male clusters, respectively, were compared. Pathways in all analyses were prioritized by the combination of the numbers of genes from the pathway involved, and the corrected p-value for the relevance of the pathway. **(B)** Calcium/calmodulin signaling was the most significantly involved shared pathway between female cluster 3 and male cluster 5. **(C)** The Integrin Signaling pathway was the most significant female specific pathway. Genes that were up- and down-regulated in fc3 compared to the other female clusters are in red and blue boxes, respectively. **(D)** Cell Cycle Regulation was the most significant male specific pathway. Genes that were up- and down-regulated in mc5 compared to the other male clusters are in red and blue boxes, respectively. See **Supplemental Table 4** for complete gene lists and statistics for each analysis.

Better outcome in mc5 was associated with The Cell Cycle Regulation pathway (**Figure 5D**). Seventeen transcripts were components of this pathway, and they included known critical regulators of mitosis such as *CDC20* (37, 38), *CKS2* (39), *PRC1* (40), *NUSAP1* (41), *PBK* (42), *Cyclin B1* and *B2* (43) and *KIF20A* (44). Fifteen of the 17 transcripts were significantly downregulated in mc5 compared to the other male clusters (p < 0.0061 for differences in the original expression data, *p* ≤ 1.7e-06 for difference in male-specific expression data) and approached the levels present in fc3 (**Figure 6A, B,** **Supplemental Figure 6**). *NEFH* and *NEFM* were the exceptions, each exhibiting greater expression in mc5 compared to the other clusters. This suggests that treatment response and survival in males is significantly determined by lower activity in factors that promote cell cycle progression.

**Figure 6:**
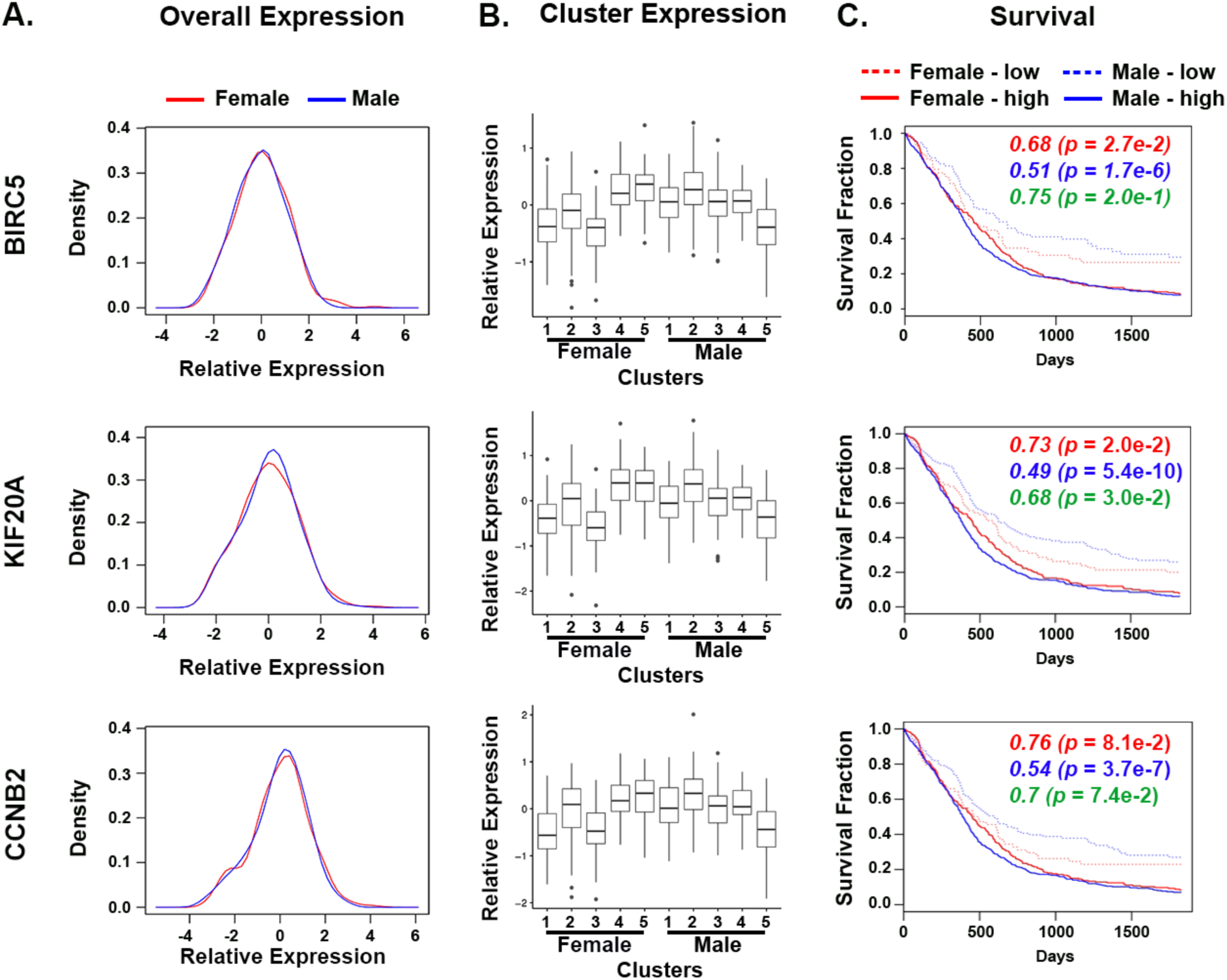
Male Cluster 5 defining genes exhibit sex-specific effects on overall survival in the merged TCGA, GSE16011 AND GSE13041 dataset. **(A)** Density plots for sex-specific expression of male (in blue) and female (in red) GBM specimens of three male cluster 5 defining genes (*BIRC5*, *KIF20A*, *CCNB2*). The overlay in male and female plots indicates near identical levels of expression in the populations. **(B)** Expression of each gene by sex and sex-specific clusters is presented as boxplots. **(C)** High and low expression groups for each gene were defined relative to the level that distinguished male cluster 5 from the other male clusters. The survival effect of differences in expression was determined for males and females. Each gene exerts a greater effect on survival in males compared to females. Cox regression hazard ratios and the corresponding p values comparing patients of low versus high expression of each of the three genes within female, within male and the interaction test comparing the effect in males to that in females are presented in red, blue and green, respectively. A parallel analysis of the other male cluster 5 defining genes is presented in **Supplemental Figure 6.**

Each of the 9 genes that distinguished fc3 (fc3.9) and the 17 genes that distinguished mc5 (mc5.17) from other female and male clusters were expressed at similar levels overall in male and female GBM patients (**Figure 6A,** **Supplemental Figure 6**). Thus, we wondered whether these genes might exert sex-specific effects on survival. For each transcript, we separated all male and female cases into low and high expression groups based on the level of expression that distinguished mc5 or fc3 from the other male or female clusters. We then determined the effect on overall survival for each transcript in each sex separately. Finally, we compared the effect of the whole gene set on overall survival between males and females in the combined dataset. None of the distinguishing genes of fc3 exhibited a differential effect on survival in males compared to females (data not shown). In contrast, while each of the downregulated cell cycle pathway genes in mc5 genes affected overall survival in both males and females, they tended to exhibit a greater effect in males compared to females (**Figure 6C,** **Supplemental Figure 6C, F**). Comparing the survival effect of the gene set in males and females, the hazard ratios of the 17 genes were significantly higher in males than their female counterparts (Wilcoxon signed rank test p=4.6e-05), indicating the gene set exerted together a significantly greater effect in males than in females, despite almost overlapping expression density of each gene in males and females (**Figure 6A**).

### Expression of sex-specific cluster defining genes correlates with chemotherapy sensitivity

Sex differences in GBM survival could result from many different cellular, tissue, or organismal factors. In order to further evaluate the potential prognostic value of the mc5.17 and fc3.9 gene signatures, we performed dose response analyses for temozolomide, etoposide, lomustine, and vincristine in five male and four female primary GBM cell lines to determine how expression levels of mc5.17 and fc3.9 specific genes correlated with IC_50_ values.

Only one cell line (male B66) demonstrated appreciable MGMT expression as measured by western blot analysis (**Supplemental Figure 7**). The temozolomide IC_50_ of this line was less than two of the other male cell lines with no MGMT expression, indicating that MGMT expression was not a dominant determinant of temozolomide resistance in these assays. Absolute IC_50_ values were calculated from each dose-response curve and were correlated with gene expression as determined by the Illumina HumanHT-12 v4 expression microarray for each cell line. Overall, male cell lines did not exhibit significantly higher absolute IC_50_ values than female cell lines (**Figure 7A**). To determine whether the levels of mc5.17 and fc3.9 gene expression stratified response for male and females cell lines, respectively, we calculated Spearman rank correlation coefficients for the relation between IC_50_ values and gene expression. Spearman rank correlation coefficients between the expression levels of the 17 genes and IC_50_s were, on average, positive for male cell lines, indicating that low expression of mc5.17 genes correlated with low IC_50_ values (high treatment efficacy) for each of the four agents, temozolomide, etoposide, lomustine, and vincristine (**Figure 7B**). In contrast, in female cell lines, low expression of the mc5.17 genes predicted high IC_50_ values (low treatment efficacy). As a negative control, the distribution of the averaged correlation coefficient of 17 randomly selected genes of 1000 random gene sets centered around 0, indicating no correlation, as expected. When the relationship between each fc3.9 gene and IC_50_ values for these drugs in male or female cell lines was analyzed, treatment efficacy in female, but not male cell lines was predicted by fc3.9 genes in 3 of the 4 drugs. Again, 1000 random gene sets of the same size of 9 randomly selected genes were not correlated with IC_50_ in any drug (**Figure 7C**). These results indicate that sex-specific expression of these genes is predictive of treatment efficacy *in vitro* and correlate with survival in patients.

**Figure 7:**
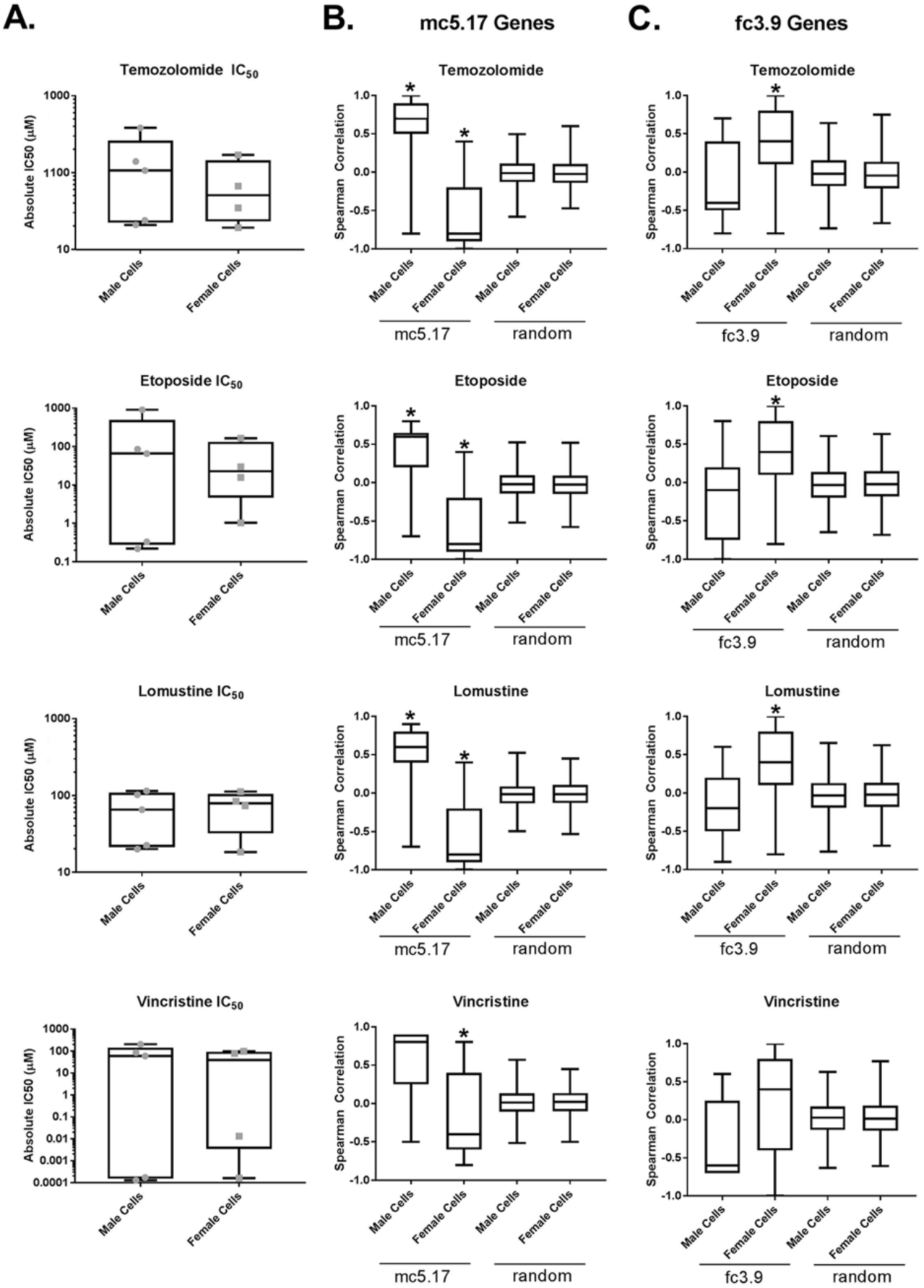
Expression of cluster defining genes correlates with sex-specific response to common chemotherapeutics *in vitro*. **(A)** Absolute IC 50 values for temozolomide, etoposide, lomustine, and vincristine for 5 male and 4 female patient-derived glioblastoma cell lines were calculated from six-point dose response curves for each cell line. Boxplots of IC_50_ across cell lines by sex were presented (horizontal bar indicates median). Median male and female IC_50_ values were not significantly different based on two sample t-test. Spearman correlation coefficient of IC_50_ values for each drug with expression of either mc5.17 genes **(B)**, fc3.9 genes **(C)**, or random gene sets in male and female cell lines. Expression of mc5.17 genes are mostly positively correlated with IC_50_ in male cell lines and negatively correlated with IC_50_ in female cell lines. Expression of fc3.9 genes are positively correlated with IC_50_ in female cell lines and not correlated with IC_50_ in male cell lines. No correlation overall existed for 1000 sets of 17 randomly selected genes **(B)** or 9 randomly selected genes **(C)**. For mc5.17 and fc3.9 genes, box plots represent the distribution of the 17 or 9 cluster-defining genes, respectively, and for random gene sets, the box plots represent the distribution of the Olkin-averaged Spearman correlation coefficient of 17 or 9 randomly selected genes per random gene set for 1000 random gene sets. Asterisks represent p<0.01 against random gene sets for each sex.

## Discussion

Sex differences are increasingly recognized as significant determinants of human health and disease. While sex differences in incidence, disease phenotype and outcome are well described and broadly recognized, the molecular bases for sex differences beyond acute hormone actions are poorly understood. Among the obstacles to improved understanding of sex differences is the inconsistent application of methodologies into lab-based and clinical research design that can adequately detect and quantify sex differences. As an example, current epidemiological data indicate that, in the United States, the male to female incidence ratio for GBM is 1.6:1 (*5*). While substantial sex differences in the incidence of glioblastoma and other brain cancers have been recognized for decades, large-scale analyses continue to most commonly merge data from both sexes, obscuring discovery of valuable information contained in the sex differences.

Recent exceptions illustrate the value of using sex differences to highlight important elements of cancer biology and clinical response. We recently found that sex-specific, cell intrinsic responses to loss of p53 function render male astrocytes more vulnerable to malignant transformation compared to female astrocytes (*17*). These findings may well relate to the sex differences in glioma incidence and are consistent with other data describing sexual dimorphism in the p53 pathway, including radiographic sex differences in men and women with glioblastoma as a function of their p53 mutational status (*37*). Understanding the molecular basis for sexual dimorphism in the p53 pathway and what it means with regard to cancer biology and clinical oncology remains an important area of research.

Most importantly, these studies emphasize that analyses without consideration of sex can obscure critical elements of biology and in aggregate, highlight the importance of parallel but separate analyses of male and female cells, male and female animals, and male and female patients. Here, we applied the JIVE algorithm to decompose male and female GBM transcriptome datasets accessed through the UCSC Cancer Genomics Browser to joint and sex-specific components. We found that male and female GBM patients cluster into five distinct male and female subtypes that are distinguished by gene expression and survival. These clusters were identified in a discovery analysis using the TCGA transcriptome dataset and validated in two independent datasets. Thus, while GBM has recently been identified as a “low sex-effect” cancer at the transcriptome level (*38*), even genes expressed at similar levels in males and females can impart substantial sex-specific effects on survival and yield mechanistically important information. Together with the sex-specific effects of p53 loss (*17*) and *Arim1* variants (*39*), these data suggest that the cellular and organismal sex context of gene expression impacts the consequences of oncogenic events. A similar mechanism was invoked to explain the sex-specific effects of AC8 polymorphisms, which elevated the risk of low grade glioma in females NF1 but lowered the LGG risk in males with NF1 (*14*).

Most compelling in this regard are the molecular features of the better survival subtypes of male and female patients. IDH1 mutation is frequently regarded as a molecular marker of better outcome in GBM (*40*, *41*). In the combined dataset, almost all IDH1 mutant tumors were assigned to fc3. This was the only female cluster with distinctly better outcome. In contrast, IDH1 mutations were distributed across all male clusters. Thus, while IDH1 mutation appears to independently influence survival in female GBM, it does not independently confer the same survival advantage in males. This finding is in contrast to a recent immunohistochemical analysis of IDH1 mutation in a single cohort of 105 patients (*42*). In this cohort, there were a total of nine IDH1 mutant tumors, 4 in males and 5 in females. The difference in survival for male patients (n=61) with and without IDH1 mutations reached statistical significance. This was not true for the female patients (n=44), but the sample size was small and the results were not validated in an independent cohort. The sex-specific impact of IDH1 mutation on survival will require additional evaluation.

The survival benefit of fc3 and mc5 was evident regardless of whether tumors were assigned to Mesenchymal, Neural, or Proneural GBM subtypes. These two clusters shared a distinguishing signature in calcium/calmodulin signaling with a particular representation of genes essential for synaptic function. They diverged in other molecular features with mc5 exhibiting a significant downregulation of mitotic spindle and cell cycle regulatory genes and fc3 exhibiting a substantial downregulation of integrin signaling pathway components. Most compelling was the sex-specific effect on survival of genes within the cell cycle regulatory pathway despite the fact that the component transcripts were expressed at similar levels in male and female tumors. These observations are consistent with the hypothesis that sex effects in cancer cannot simply be defined by the levels of gene expression, but rather need to include the potential sex differences in gene effect. A similar observation regarding sex differences in the significance of MGMT promoter methylation was recently published (*42*). In three independent cohorts of patients, MGMT promoter methylation was shown to impact on survival of female, but not male GBM patients. Thus, moving forward, investigations into the mechanisms of sex differences in biology should focus more on sex-specific effects of genes rather than levels of gene expression.

Among the striking results of this study is the ready harmonization of the effect of sex on gene expression, the *in vitro* drug sensitivity, and the MRI measures of tumor dispersion and proliferation. The gene expression analysis identified downregulation of cell cycle progression and downregulation of integrin signaling as significantly correlated with best survival in male and female patients, respectively. Expression levels of the 17 and 9 gene signatures that distinguished the longest surviving male and female cohorts, respectively, was also shown to directly correlate, in a sex-specific manner, with *in vitro* drug sensitivity as measured by IC_50_ values, for a panel of primary GBM cell lines. Moreover, we found evidence that MRI-based predictors of survival may differ for males and females with GBM. These predictors are based on measures of rates of proliferation and invasion. Although the biological mechanisms underlying these MRI-derived measures of net tumor proliferation and invasion are complex, the observed phenotypic differences in the MRI analysis are consistent with the gene expression analysis. The decreased net proliferation rate (ρ) observed in long-lived males is consistent with the pathway enrichment for cell-cycle regulation identified by the gene expression analysis of long-lived mc5. Further, the increased net invasion rate (D) in the long-lived female patients is consistent with the pathway enrichment for integrin signaling observed in long-lived fc3.

While there was no statistical significance between survival for the longest-lived male and female patients, these data suggest that survival is determined by different factors for males and females with GBM. The MRI analysis suggests that while long-lived female patients tend to have more aggressive tumors both in terms of net proliferation (ρ) and net invasion (D) rates prior to surgery, these female GBM patients receive greater benefit from treatment as evidenced by decreased tumor growth velocity during temozolomide treatment. This suggests that treatment, rather than innate tumor biology may be the dominant determinant of survival in females. Conversely, long-lived male patients tend to have less aggressive tumors prior to surgery with smaller D and ρ, and exhibit little to no change in tumor growth velocity with temozolomide. This suggests that long-lived male patients may achieve their survival less from their response to treatment and more from innate tumor biology.

Finally, the current study suggests that greater precision in GBM patient stratification may be achieved with this approach of sex-specific molecular subtyping and that improvements in GBM outcome might be possible with sex-specific approaches to treatment, including blocking cell cycle progression in male patients and targeting integrin signaling in female patients. The value of the male proliferative and female cell adhesion phenotypes should be further evaluated in additional datasets, including recent clinical trial results of cell cycle inhibitors and integrin antagonists, as well as in prospective studies.

## Materials and Methods

### Patients and Samples

Primary human GBM specimens for culture were obtained and utilized in accordance with a Washington University Institutional Review Board (IRB)-approved Human Studies Protocol (#201102299). Expression profiling of total RNA extracted from each GBM cell line was performed in replicate for each line using the Illumina HumanHT-12 v4 expression microarray platform. Patient clinical treatment and related imaging data were obtained in accordance with a Mayo Clinic IRB-approved Human Subjects Protocol (#15-002337).

### MRI Image Analysis

The clinical research database of the Mayo Clinic (Phoenix) was searched for newly diagnosed GBMs patients who 1) received standard-of-care maximum possible resection, post-operative radiation therapy and temozolomide (TMZ) chemotherapy (*21*) and 2) with sufficient imaging to estimate patient-specific net invasion rate (D), and net proliferation rate (rho (ρ)). A total of 111 patients, 74 males and 37 females met these criteria. For each patient, we segmented the T1Gd and T2 MRIs to calculate lesion volumes using a house-built method in Python using standard image segmentation algorithms. Volumes (V) were converted into spherically equivalent mean radii, r= (V/4π)∧(1/3). Serial imaging during adjuvant temozolomide allowed estimation of the tumor growth velocity between images obtained at 2 month intervals, velocity = (r_t2-r_t1)/(t2-t 1). Further, following established methods (*23*, *25*, *28*, *43*-*46*), tumor radii on T1Gd and T2 MR images were then used to estimate the patient-specific net invasion rate, D, and net proliferation rate, ρ. Specifically, we estimated D and ρ using methods attuned to patients with only 1 MRI time point prior to treatment (*28*). This approach utilizes the size ratio of necrosis to the entire tumor as a surrogate marker of aggressive growth or velocity. This is implemented via a combination of the proliferation-invasion model and a second mathematical model which decomposes the tumor population into three phenotypes: normoxic, hypoxic, and necrotic. This second model is called the Proliferation-Invasion-Hypoxic-Necrotic-Angiogenesis model, or the PIHNA model, and also includes D and ρ as critical model parameters (*43*). A lookup table was created from numerous runs of the PIHNA model, each distinguished by different values of D and ρ, with other parameters remaining constant. Analysis of simulations of the PIHNA model for various D and ρ (with all other parameters constant) reveals a relationship between the size of necrosis, and the size of the bulk tumor (>=80% tumor) for a given size of the invading tumor (>=16% tumor). Given a patient’s measurements from a single time point of the T1Gd MRI (associated with >=80% tumor), the T2 MRI (associated with >=16% tumor), the necrosis, and the PI estimated D/ρ, a D and ρ values are returned.

### Transcriptome Data

The processed whole-transcriptome gene expression data of 539 GBM patients profiled on the Affymetrix HT Human Genome U133a microarray platform, and related phenotypic information (sex, survival, TCGA molecular subtypes) were downloaded from the UCSC Cancer Genomics Browser (*47*). A total of 320 male and 205 female GBM patients with complete phenotypic information were analyzed. Among 12042 genes, 8720 genes were retained for analyses based on coefficient of variation ≥0.4, **Supplemental Figure 8**). The processed whole-transcriptome gene expression data of two public data sets (GSE16011 (*48*) and GSE13041 (*49*)) were downloaded from Gene Expression Omnibus database (http://www.ncbi.nlm.nih.gov/geo). GSE16011 consists of 276 GBM patients (184 males and 92 females) profiled on Affymetrix human genome U133 plus 2.0. GSE13041 consists of 174 GBM patients (105 males and 69 females) profiled on Affymetrix human genome U133A and U133 plus 2.0 (samples profiled on Affymetrix human genomeU95Av2 were excluded due to few overlapping genes).

### JIVE Analysis

The workflow for the application of the Joint and Individual Variance Explained (JIVE) analysis is illustrated in **Supplemental Figure 9**. For the TCGA GBM data set, each gene was separately centered by the overall mean across all patients of both sexes. The centered data were split by sex into a male expression data set and a female expression data set. Both data sets were input to the R package “r.jive” (version 1.2), for an integrative Joint and Individual Variance Explained (JIVE) analysis. JIVE identifies dominant expression patterns across samples like the principal component analysis (PCA), but with the additional ability to attribute expression variations to shared and individual patterns of both sexes, and thus, to derive the joint expression component shared by both sexes, the individual sex-specific expression component and other residual components (*50*).

### Definition of Sex-specific Clusters

To define sex-specific subtypes, the hierarchical clustering method was applied to each sex-specific expression component to group patients into subsets of relatively similar expression patterns according to the distance metric of one minus Pearson correlation using the average linkage. In order to determine the number of clusters and identify “core” samples belonging to each cluster, the resampling based consensus clustering method was performed using the R package “ConsensusClusterPlus” (*51*). Two types of consensus clustering were performed, an un-weighted one, and one with samples weighted by the explained variation of the sex-specific components. Samples were considered as uncertain and removed if they were grouped differently in the two types of clustering, or if they had negative silhouette scores, which measures how similar a sample in one cluster is to others in the same cluster (*52*). This affected 65 of the 205 female and 100 of the 320 male samples. Small clusters composed of too few samples (<10) were also removed. The number of clusters was determined by the delta area under the consensus cumulative distribution function (CDF). The area under the CDF and the delta change (relative change from k to k+1 clusters) were used to quantify the concentration of the consensus distribution using ConsensusClusterPlus (*51*, *53*). Based on the delta area plots from ConsensusClusterPlus, the consensus increase became insubstantial from 5 clusters to 6 clusters (delta ~=0.05) in both males and females, using either unweighted or weighted clustering.

### Signature Genes and Pathway Analysis

To identify cluster-defining gene signatures, differential gene expression analysis was performed using the significance analysis of microarray (SAM) method implemented in the R package “siggenes” (*54*) to compare one sex-specific cluster versus the other four sex-specific clusters. Genes showing significant expression differences at the 5% false discovery rate (FDR) were identified as the cluster defining signature genes. The official gene symbols of genes differentiating each cluster from the remaining clusters within a sex were imported into the Genomatix Pathway System (GePS, http://www.genomatix.de) for enriched biological pathways and GO terms. FDR adjusted *P*-Values were reported.

### Survival Analysis

Overall survival and disease free survival were the primary endpoints for survival analyses. The Kaplan-Meier (KM) product limit method was used to estimate the empirical survival probabilities, and survival differences among groups were compared by the log rank test using the *R* package “survival” (*55*). The raw hazard ratios between groups were estimated as the ratio of their relative event rates calculated from the ratio of observed to expected events (*56*). The TCGA DFS and OS were censored up to 5 years in consideration for robust statistical analyses.

### Overall Sex-Specific Survival Effect of the Male Cluster 5 Gene Set

Pathway analysis identified Cell Cycle Regulation as the top enriched pathway in the best surviving male cluster (mc5). To investigate the overall sex-specific effect of the 16 cluster-defining genes on male and female survival, we first dichotomized the continuous gene expression. For each gene, a cutoff value of the gene’s expression in the extracted male-specific expression components was identified such that the sum of classification error rates was minimized between the mc5 versus the other male clusters. Both male and female patients were subsequently divided into low and high expression group by the cutoff value corresponding to each gene. The hazard ratios (HRs) of the 16 genes (high versus low expression) in males and females were separately estimated from the Cox proportional hazard model to measure their effect on male and female survival. One-sided Wilcoxon signed rank test was used to compare the HRs between male and female to gauge the significance of the overall sex-specific effect of the gene set on survival and test the specific assumption that the effect would be greater in males compared to females.

### Result Validation

We used the two independent datasets (GSE16011 and GSE13041) to validate the survival difference between the sex-specific clusters defined using the TCGA prototype data. The two datasets were merged with the TCGA samples using the COMBAT method (*57*) implemented in the R package “inSilicoMerging” (*58*). The expression of the merged data was then reversely linearly transformed so that the TCGA expression data remained the same before and after merging, and the validation samples had expression levels that were similar in scale to the TCGA samples. Validation samples were assigned to TCGA molecular subtypes by the 1-nearest-neighbor method based on the Euclidian distance metric. The transformed validation data were separated by sex and the joint and sex-specific expression were derived by projecting onto the JIVE principal components of the TCGA samples. Based on the extracted sex-specific expression and Pearson correlation coefficient, each male and female sample in the validation sets was assigned to the closest male and female clusters among all sex-specific clusters defined with the TCGA samples. Survival curves of the resulting sex-specific clusters were generated by the KM method and survival differences were assessed by log-rank test and hazard ratios from Cox model.

### In vitro drug screens

Resected human GBM resection tissue was mechanically dissociated with Accutase (Sigma) and cultured on Primaria plates (BD Biosciences) coated with laminin (Sigma) with RHB-A media (Takara) supplemented with EGF (Sigma) and bFGF (Millipore) (*59*). Dose response curves were performed in a 384-well plate format (400 cells/laminin-coated well in triplicate). Cells were treated for 48 hours (lomustine, vincristine) or 72 hours (etoposide, temozolomide) at various doses, and Cell Titer Glo (Promega) was used to assess cell number (relative to vehicle treated control). Absolute IC_50_ values were determined in GraphPad Prism by interpolating results from standard curves derived from the least-squares fit of data from three separate experiments.

### Western blot analysis

Primary mes-GBM stem cell cultures were lysed in RIPA buffer in the presence of complete protease inhibitor (Sigma). Thirty μg of cleared protein was separated on a 4-12% Bis-Tris acrylamide gel and transferred to nitrocellulose. After blocking with Odyssey Blocking Buffer (Licor) overnight at 4°C, blots were incubated with primary antibodies to actin (Sigma, 1:40,000) and MGMT (RD Biosystems, 1:200) overnight at 4°C, followed by incubation with secondary antibodies (Licor), and imaged using the Odyssey Imaging platform.

## Acknowledgements

This work was supported by grants from the NIH, R01 CA174737 (JBR), R01 NS060752 (KRS), R01 CA164371 (KRS), U54 CA210180 (KRS), U54 CA143970 (KRS), U54 CA193489(KRS), NIH K08 NS081105 (AHK), NIH R01 NS094670 (AHK). The Children’s Discovery Institute of Washington University (JBR), Joshua’s Great Things (JBR). The James S. McDonnell Foundation, the Ivy Foundation, and the Mayo Clinic (KRS). NIH U01 CA168397 (MEB), Ben & Catherine Ivy Foundation (MEB). The content is solely the responsibility of the authors and does not necessarily represent the official views of the National Institutes of Health or Government.

## Supplementary Materials

### Supplemental Figures

Supplemental Figure 1: Geometric illustration of JIVE decomposition.

Supplemental Figure 2: Identification of the sex-specific clusters by consensus clustering.

Supplemental Figure 3: The joint structure of the male and female TCGA GBM samples captures the dominant molecular signatures of TCGA subtypes.

Supplemental Figure 4: Expression variation in TCGA data explained by the JIVE components.

Supplemental Figure 5: Overall survival analysis of sex-specific clusters in TCGA data.

Supplemental Figure 6: Male cluster 5 defining genes exhibit sex-specific effects on survival.

Supplemental Figure 7: Western blot analysis of MGMT expression in female and male patient-derived glioblastoma cell lines.

Supplemental Figure 8: Distribution of the coefficients of variation in the TCGA data.

Supplemental Figure 9: Analysis flowchart.

### Supplemental Tables

Supplemental Table 1: Gene expression signatures for sex-specific clusters.

Supplemental Table 2: Distribution of Verhaak subtypes to sex-specific clusters

Supplemental Table 3: p-values and hazard ratios associated with survival differences in sex-specific clusters.

Supplemental Table 4: Pathway analysis of female cluster 3 and male cluster 5.

